# Identifying differences in gait adaptability across various speeds using movement synergy analysis

**DOI:** 10.1101/2020.07.15.203307

**Authors:** David Ó’Reilly, Peter Federolf

## Abstract

**Introduction:** The aim of this study was to identify movement synergies during normal-walking that can differentiate healthy adults in terms of gait adaptability at various speeds. To this end, the association between movement synergies and lower-limb coordination variability or Deviation Phase (DP) was investigated. A secondary aim of this study included an investigation into the moderating effect of these movement synergies on the relationship between DP and the smoothness of arm-swing motion quantified as the normalised jerk index (NJI).

**Method:** A principal component analysis of whole-body marker trajectories from normal-walking treadmill trials at 0.8m/s, 1.2m/s and 1.6m/s was undertaken. Both DP and NJI were derived from approx. 8 minutes of perturbed-walking treadmill trials. Principal movement components, PM_k_, were derived and the RMS of the 2^nd^-order differentiation of these PM_k_ (PA_k_RMS) were included as independent variables representing the magnitude of neuromuscular control in each PM_k_. The PA_k_RMS were input into separate maximal linear mixed-effects regression models to explain the variance in DP and (DP × NJI). A stepwise elimination of terms and comparison of models using Anova identified optimal models for both aims.

**Results:** Among the first 7 validated PM_k_, PA_4_RMS (double-support phase) was identified as an optimal model and demonstrated a significant negative effect on DP however this effect may differ considerably across walking-speeds. An optimal model for describing the variance in (DP × NJI) included a fixed-effect of PA_6_RMS (Left – Right side weight transfer). Within-participant clustering was prevalent within both optimal models.

**Interpretation:** The hypotheses that individuals who exhibited greater control on specific kinematic synergies would exhibit variations during perturbed walking was substantiated. Supporting evidence for the role of movement synergies during the double-support phase of gait in proactively correcting balance was presented. The potential influence of leg dominance on gait adaptability was also discussed. Future studies should investigate further the role of walking-speed and leg dominance on movement synergies and look to generalize these findings to patient populations.

**Highlights:**

- Baseline movement synergies representing terminal-swing and double-support phases of gait were found to have significant negative effects on lower-limb coordination variability during perturbed-walking trials at various speeds.
- Movement synergies related to the double-support phase and weight transfer events of gait were determined to have a negative moderating effect on the translation of lower-limb coordination variability into upper-limb postural corrections.
- Evidence was presented for the important role of the double-stance phase of gait in gait adaptability while leg dominance was shown to play a potential role in differentiating healthy adults in this study.

## 1.1. Introduction

Stability and adaptability of human movement during quiet stance and locomotion is an area of increasing relevance in the growing aging population. Approximately 30% of older adults will experience a fall leading to extensive personal, social and economic costs (Davis *et al.*, 2010; Soriano, DeCherrie, & Thomas, 2007). A stable gait pattern can be achieved when the effects of small perturbations throughout each stride can be limited, when balance can be recovered successfully from large perturbations and by not encountering perturbations in the environment that exceed the limits of the system (Bruijn *et al.*, 2013). To prevent small perturbations at each stride from accumulating into a serious instability, human walking takes on an inverted pendulum like motion that effectively harnesses the biomechanical constraints of the lower-limbs to allow for smooth step-to-step transitions that are both energy efficient and adequately stable (Kuo, 2007). When required, postural control strategies are also introduced by the central nervous system (CNS) to accommodate fall prevention. These may include a rapid change in swing-foot placement trajectory and modulation of stance-foot plantar-flexion at propulsion phase (Reimann *et al.*, 2018; Young & Dingwell, 2012). This may also be coupled with a change in arm-swing motion towards a more elevated and lateral position to adjust the bodys center-of-mass (Inkol, Huntley, & Vallis, 2019). Inkol, Huntley & Vallis (2019) noted that the magnitude and direction of such reactive arm-swing strategies varied considerably across healthy participants despite a uniform perturbation being adminstered.

To ensure stability while also achieving adaptability, an optimal level of variability in the dynamics of gait is necessary (Hausdorff, 2005). This trade-off is a phenomenon that is present in various physiological signals (e.g. heart-rate variability) (Stergiou & Decker, 2011). Variability above or below this optimal level can be indicative of pathology as excessive periodicity causes a system to be insufficiently adaptable to the demands of the environment while excessive variability introduces sub-optimal performance and instability (Hausdorff, 2005; Manor *et al.*, 2010). Evidence from studies investigating this phenomenon through the dynamic systems perspective suggests that abnormal gait coordination variability is related to instability in patient populations (Hacmon *et al.*, 2012; Paterson, Hill, & Lythgo, 2011). Nonetheless among healthy adults, both high- and low-variability can be associated with gait stability indicating that other factors may also be at play in producing instable gait (Beauchet *et al.*, 2009). Variability and changes in variability may also reflect the adaptability of gait and its adaptions to environmental demands (Madehkhaksar *et al.*, 2018). Therefore, the context and subject in question must be considered when assessing movement from this perspective.

Human movement is thought to be modularly controlled by the CNS through task-relevant synergistic muscle activations and in doing so, the CNS selects a near optimal solution to the motor-task from the many redundant degrees of freedom available (Berret *et al.*, 2019; Bizzi & Cheung, 2013). Common, invariant muscle synergies have been identified across various activities such as standing and walking (Chvatal & Ting, 2013) while task-specific synergies have also been revealed during slipping for instance (Nazifi *et al.*, 2017) demonstrating the adaptability of human motor control to environmental demands. Nazifi et al. (2017) revealed differences across healthy adults in terms of slip-severity were related to the magnitude of specific muscle activations in response to a perturbation (e.g. medial hamstring). In a similar vein, Robert et al. (2009) demonstrated the differential control the CNS exhibits on body segments in maintaining whole-body angular momentum during the swing-phase compared to the double-support phase of gait. Analysis of muscle/movement synergies remains an under researched field with the potential to provide a generalisable methodology for assessing balance performance across the gait-cycle (Chvatal & Ting, 2013).

The measurement of discrete points in complex systems requires their synthesis to give more accurate and reliable interpretations. The *‘curse of dimensionality’* plays a major role in this process as large feature sets are to be analysed simultaneously. Dimensionality reduction methods have become popular in this the era of *‘big data’* as they alleviate this issue by reducing the number of features needed for a comprehensive analysis (Phinyomark *et al.*, 2018). This reduction can be implemented via the creation of a smaller number of new variables that retain a large amount of the information in the original feature-set. Alternatively, through feature selection methods the most relevant feature-subset can be identified and extracted based on specific criteria. Dimensionality reduction has been used successfully to identify muscle synergies (Steele, Rozumalski, & Schwartz, 2015), contrast sporting techniques (P. Federolf *et al.*, 2014), improve fall-risk classification (O’Reilly, 2019; Rocchi, Chiari, & Cappello, 2004) etc. and their uses continue to expand with the availability of large datasets.

In particular and of specific relevance to this study, recent research has found effective use of principle component analysis (PCA) in reducing the dimensionality of kinematic data for various aims (Daffertshofer *et al.*, 2004; Federolf, 2016; Troje, 2002). Of note, the ‘*Principle movements’* (PM) are the eigen-vectors derived from a PCA of movement data that represent correlated changes in marker coordinates (synergies) (Federolf, 2016). Each successive PM extracted from this analysis is ordered in terms of the amount of variance explained by the movement with higher-order PMs explaining less variance and representing more subtle movement patterns (Phinyomark *et al.*, 2015). It is of interest in this study to what extent the neuromuscular system controls each of the PM_k_ components identified in subjects. By projecting the PM_k_ scores onto a posture space, the change in position of the body segments can be represented with respect to time and is known as the ‘*principal position’* (PP_k_) (Federolf, 2016). In accordance with Newton’s laws of motion, PP_k_ time-series can be differentiated to ‘*principal velocities’* (PV_k_) and ‘*principal accelerations*’ (PA_k_) representing the 1^st^- and 2^nd^-order derivatives of the PP_k_ time-series (Federolf, 2016). Variables computed from PA_k_ time-series have been of focus in some of the related literature as the extracted accelerations are thought to represent the actions of the neuromuscular system on body segments (Haid *et al.*, 2018; Promsri, Haid, & Federolf, 2020; Zago *et al.*, 2019).

In this study, a novel PA_k_ variable (PA_k_RMS) representing the magnitude of neuromuscular control on PM_k_ will be derived from a gait analysis of healthy adults exposed to mechanical perturbations at various speeds. The aim of this study is to determine the utility of PA_k_RMS in identifying key gait mechanisms that differentiate healthy individuals in terms of gait adaptability. To this end, the following associations will be investigated:

1) PA_k_RMS and lower-limb coordination variability and 2) the moderating effect of PA_k_RMS on the relationship between lower-limb coordination variability and smoothness of arm-swing motion. It is hypothesized for the first aim that PA_k_RMS will demonstrate a negative effect on lower-limb coordination variability as participants who exhibit more control on specific PM_k_ will experience less disturbance in their natural flow of walking and therefore require less adaptations. For the secondary aim, it is hypothesized that PA_k_RMS will demonstrate a negative moderating effect on this upper-limb – lower-limb relationship as individuals who exhibit greater control of specific PM_k_ will also be able to prevent lower-limb perturbations from accumulating into trunk-level postural corrections.

## 2.1 Methods

### 2.1.1 Secondary Data analysis

Thirty-nine trials of normal and perturbed treadmill walking collected from 13 participants were taken from an open-source dataset generated in the Human Motion and Control Laboratory in Cleveland State University (J. K. Moore, Hnat, & van den Bogert, 2015). Characteristics of the included participants were: Age (Years)= 24±4.12, Height (Meters)= 1.74±0.08, Weight (kg)= 72.88±12. In short, participants were asked to walk normally on an R-Mill treadmill with dual 6-degrees of freedom force plates and independent belts for each foot (Forcelink, Culemborg, The Netherlands). Motion capture was undertaken using a 10 Ospreys camera motion capture system paired with the Cortex 3.1.1.1290 software at a sample rate of 100Hz (Motion Analysis, Santa Rosa, CA, USA). Participants were first asked to walk for 2 minutes unperturbed and were then subjected to 8 minutes of longitudinal belt-speed perturbations. This protocol was repeated for each of 0.8m/s, 1.2m/s and 1.6m/s walking speeds. The pseudo-random belt-speed control signals used to induce perturbations during each stance phase were generated a priori using MATLAB and Simulink (Mathworks, Natick, Massachusetts, USA) and are available online (Hnat, Moore, & Van den Bogert, 2015).

Marker trajectories from a 47 whole-body marker setup for both normal-walking and perturbed-walking trials were extracted and processed using the GaitAnalysis toolbox V 0.1.2 (J. Moore *et al.*, 2014). Data for normal-walking trials was taken from when a constant belt-speed was achieved (leaving approximately 50 seconds of normal-walking) and the length of the time-series were normalized to 5,000 data points and then low-pass filtered with a 4^th^-order Butterworth filter at a cut-off frequency of 20Hz.

### 2.1.2 Independent variable computation

All of the beforementioned procedures in this section were carried out using PManalyzer software (Haid *et al.*, 2019), a Matlab GUI specifically designed for PM variable computation (Matlab (R2019B), Natick, Massachusetts: The MathWorks Inc.). Only data from normal-walking trials was included in the independent variable computation. In short and in line with other recent endeavours using this approach (Longo *et al.*, 2019; Promsri, Haid, & Federolf, 2020; Zago *et al.*, 2019), the data from all subjects and walking-speed trials were pooled into one matrix to allow for direct comparisons between subjects and across trials. Raw marker coordinates (XYZ) were first transposed so that each frame represents a posture vector, then centred by subtracting each posture vector from the mean posture vector to get postural deviations and normalized to their mean Euclidean distances to ensure an equal contribution by each subject/trial to the pooled dataset and subject-specific overall variance respectively. The data was also centred towards the centre-of-mass to create a body-position dependent coordinate system thus removing the potential inclusion of irrelevant body displacements from the PMs (P. Federolf *et al.*, 2014). The pooled dataset of concatenated normal-walking trials accumulated into a 195,000 × 60 input matrix (100 Hz [Sampling rate] × approximately 50 seconds [Trial duration] × 3 [Three trials] × 13 [Number of subjects] × 60 [Marker coordinates]). The PCA algorithm decomposed this input into a covariance matrix of PM_k_.

The PM_k_ within the output of this process were leave-one-out cross validated. The first 7 PMs presented a change of less than 15° when a participant was left out and were therefore included in further analysis (Haid *et al.*, 2018). To compute the PA_k_ time-series, the PP_k_ time-series were first extracted as the PCA-scores derived from the PM eigen-vectors that represent deviations in posture across the orthonormal posture-space (Federolf, 2016). The time-series were then further low-pass filtered using a 3^rd^-order Butterworth filter at a cut-off of 10Hz and their spectral properties inspected with a Fourier analysis for frequency power above what is expected in noise free movement data (Promsri *et al.*, 2020). No significant power was found at frequencies above 5Hz and so the effect of noise was deemed to have been sufficiently reduced. The control of PM_k_ was represented as the root-mean-square (*RMS*) of the PA_k_ time-series with respect to trial duration (*t*) (*PA_k_RMS*) (Equation 1). In previous studies using this measure, static balance tasks were investigated where the accelerations in PA_k_ components could be directly related to postural control mechanisms (Promsri, Haid, & Federolf, 2020). During dynamic movements however, it must be noted that these accelerations additionally include dependencies such as walking-speed and so cannot solely reflect the actions of the neuromuscular control system (Zago *et al.*, 2019). Therefore, the *PA_k_RMS* were then normalized by the respective walking-speed of the trial.

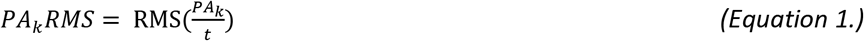

### 2.1.3 Dependent variable computation

Lower-limb coordination variability was represented as the stride-to-stride standard deviation in inter-joint continuous relative phase (CRP) values known as the deviation phase (DP). Heel-strikes events during perturbed walking trials were identified using a coordinate-based treadmill algorithm (Zeni Jr, Richards, & Higginson, 2008). CRP was calculated in accordance with the methodology described in (Hamill *et al.*, 1999) to determine the coordinative relationship between the Hip-Ankle, Hip-Knee and Knee-Ankle of both limbs in the sagittal plane only. Equation 3 below illustrates the CRP formula where φ_*proximal*_ (*t*) is the phase normalized joint angle of the proximal segment at timepoint t and φ_*distal*_ (*t*) is the phase normalized joint angle for the distal segment at the same timepoint. The average DP across the CRP measurements within each perturbed walking-speed trial was then taken as a summary statistic for further analysis.

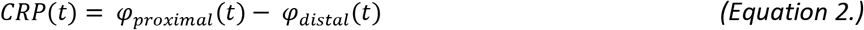

Arm-elevation jerk (*Jerk_v_*) was computed as the 3^rd^-order derivative of the medial-wrist marker position trajectories 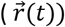 with respect to time (Equation 3). The mean absolute *Jerk_v_* 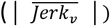 was then normalised by the peak velocity 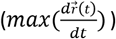 of the marker trajectory to formulate a measure of movement smoothness adopted from Roher et al. (2002) hereafter referred to as the normalised jerk index (*NJI*). As reactive arm-swing strategies are said to be asymmetrical (Inkol, Huntley, & Vallis, 2019), the side that exhibited the highest NJI value was included in statistical analyses.

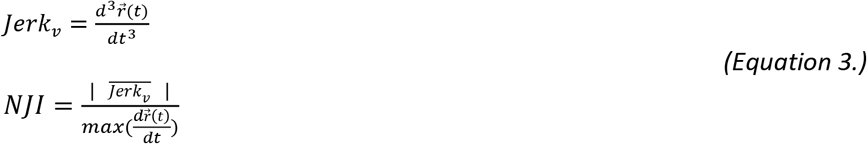

### 2.1.3 Statistical analysis

Due to the repeated-measures structure of this study along with the hypotheses being tested, a linear mixed-effects regression was deemed appropriate. Mixed-effects regression is different to the traditional ordinary least-squares in that it can flexibly consider effects at different levels of analysis (i.e. multi-level regression), known as random-effects. Mixed-effects models are useful for data structures which may have clustering present, providing more accurate effect estimates than traditional techniques in such situations (Magezi, 2015). In the current study, clustering may occur as a result of multiple observations from the same participant while the extent to which the effect of PA_k_RMS varies across trials is also of interest to this study’s aims. Barr et al. (2013) suggested that data-driven maximal models should be utilised in the case of confirmatory hypothesis modelling. Therefore, the following effect terms were included in the models for both aims:

#### PA_k_RMS

Each PA_k_RMS component will be input into separate models as fixed-effects, to examine how the magnitude of neuromuscular control on each movement synergy affects gait adaptability.

#### Trial

As gait variability is said to be speed-dependent (H. G. Kang & Dingwell, 2008), the fixed-effect of walking-speed was included in both models.

#### PA_k_RMS * Trial

It is within the scope of this study’s aims to examine the influence of walking-speed on the predictability of PA_k_RMS on the dependent variables. Therefore, an interaction term was modelled in both aims.

##### (PA_k_RMS|Participants)

As a random sample of participants were taken from the population and dependency across observations is present due to repeated measures, subject-specific intercepts were modelled allowing within-participant clustering to be considered. This was accompanied by random slopes to allow the slope to vary across participants also.

##### (PA_k_RMS |Trial)

As the walking-speeds in each trial were arbitrarily selected, the effect of this random selection on the data structure will be considered by the inclusion of a random trial intercepts term. Along with this random slopes for trials will also be included to allow for differences in the effect of PA_k_RMS across trials to be considered.

Equation 4 & 5 below illustrate the maximal models for each aim in R software syntax form. Due to the small sample size included in this analysis, a step-wise elimination of fixed and random-effect terms from these maximal models was conducted to improve parsimony. Each PA_k_RMS was separately input into individual mixed-effects models with the above described terms and a step-wise elimination was carried out for each of the 7 models in both hypotheses. The output from this procedure was then compared in Anova with F-test p-values and Satterthwaite’s degrees-of-freedom approximation to identify optimal models for both hypotheses. An optimal synergy was identified as it was likely that a number of synergies would be relevant to the dependent variables due to inherent interdependencies and reciprocal compensation rather than through a direct relationship (Fettrow *et al.*, 2019; Latash, Scholz, & Schöner, 2002). To reduce familywise error accumulation from multiple comparisons, a Bonferroni correction was implemented in which a significance level of p<0.007 was set a priori for the final models. These statistical procedures were conducted using the *‘LmerTest’* package in R software (Kuznetsova, Brockhoff, & Christensen, 2017). Goodness-of-fit was assessed using Akaike information criterion (AIC), Bayesian’s information criterion (BIC) and McFadden’s R-squared while coefficient estimates and p-values were obtained using restricted maximum-likelihood and the Satterthwaite’s degrees of freedom method respectively.

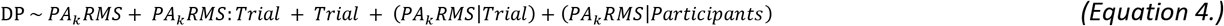

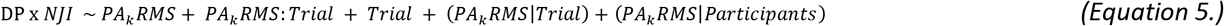

## 3.1 Results

### 3.1.1 Walking condition variables

Average and standard deviation values for the variables of interest in this study are presented in Table 1 while a summary of each validated PM_k_ is described in Figure 1. These graphical representations are also available as 2D and 3D videos in the supplementary material attached. DP consistently increased with walking-speed in the perturbed-walking trials from a low of 25.1±3.5 at 0.8m/s to a high of 28.6±2.4 at 1.6m/s. Of note however, the 0.8m/s trials experienced the highest between-participant variability in DP. The smoothness of arm-swing elevation during perturbed-walking trials (NJI) also increased with walking-speed from 0.16±0.07 at 0.8m/s to 0.24±0.14 at 1.6m/s. The non-normalized values for the first 7 validated PA_k_RMS components taken from normal-walking trials are also presented. All PA_k_RMS components demonstrated a positive relationship with walking-speed except in the case of PA_2_RMS (representing the foot placement mechanism) which exhibited a negative relationship with walking-speed. The dominant movement captured using PCA was PM_1_ (variance explained= 94.2%) that was interpreted to represent a stiff inverted-pendulum motion of the lower-limbs along with elbow-flexion. PM_3_ captured the synchronous extension of the knees during mid-sing/stance phase while PM_4_ captured the push-off mechanism. PM_5_ and PM_6_ represent step-to-step transitions from the left – right side and right – left side respectively. PM_7_ explained just 0.44% of the total variance and so a recognizable gait pattern could not be interpreted. It could be noted however that transverse plane rotations of the lower-limbs and elbows were present for PM_7_.

**Table 1:**
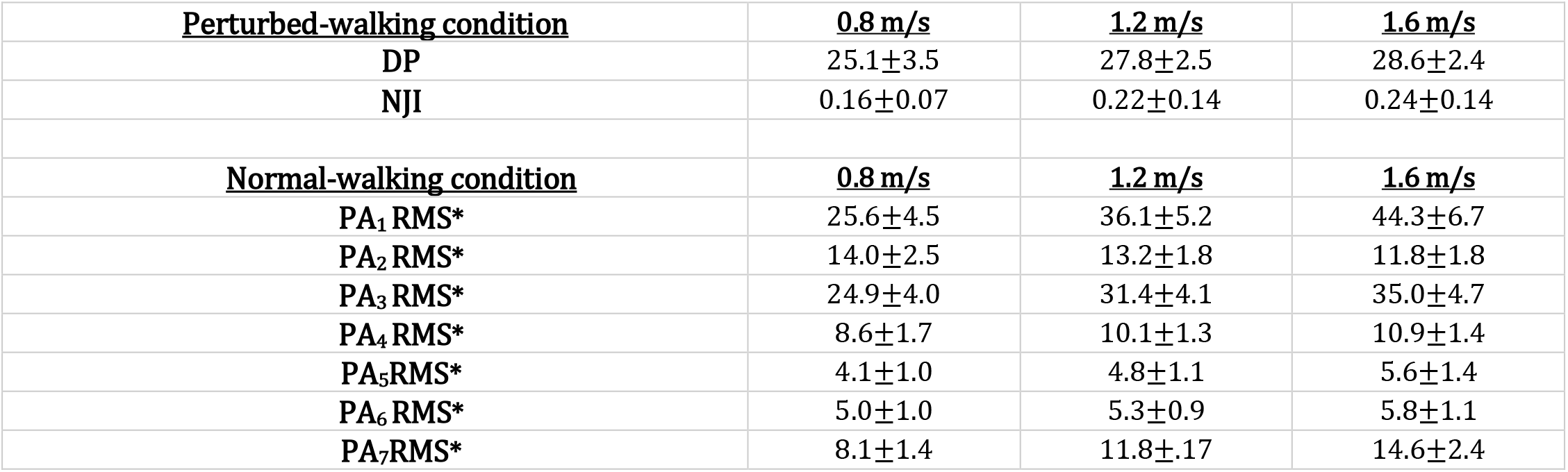
Mean and standard deviation for independent and dependent variables in both conditions and for each walking-speed. (* non-normalized values for PA_k_RMS).

| **<u>Perturbed-walking condition</u>** | **<u>0.8 m/s</u>** | **<u>1.2 m/s</u>** | **<u>1.6 m/s</u>** |
| --- | --- | --- | --- |
| **DP** | $25.1 \pm 3.5$ | $27.8 \pm 2.5$ | $28.6 \pm 2.4$ |
| **NJI** | $0.16 \pm 0.07$ | $0.22 \pm 0.14$ | $0.24 \pm 0.14$ |
| **<u>Normal-walking condition</u>** | **<u>0.8 m/s</u>** | **<u>1.2 m/s</u>** | **<u>1.6 m/s</u>** |
| **$PA_1RMS^*$** | $25.6 \pm 4.5$ | $36.1 \pm 5.2$ | $44.3 \pm 6.7$ |
| **$PA_2RMS^*$** | $14.0 \pm 2.5$ | $13.2 \pm 1.8$ | $11.8 \pm 1.8$ |
| **$PA_3RMS^*$** | $24.9 \pm 4.0$ | $31.4 \pm 4.1$ | $35.0 \pm 4.7$ |
| **$PA_4RMS^*$** | $8.6 \pm 1.7$ | $10.1 \pm 1.3$ | $10.9 \pm 1.4$ |
| **$PA_5RMS^*$** | $4.1 \pm 1.0$ | $4.8 \pm 1.1$ | $5.6 \pm 1.4$ |
| **$PA_6RMS^*$** | $5.0 \pm 1.0$ | $5.3 \pm 0.9$ | $5.8 \pm 1.1$ |
| **$PA_7RMS^*$** | $8.1 \pm 1.4$ | $11.8 \pm 1.7$ | $14.6 \pm 2.4$ |

**Figure 1:**
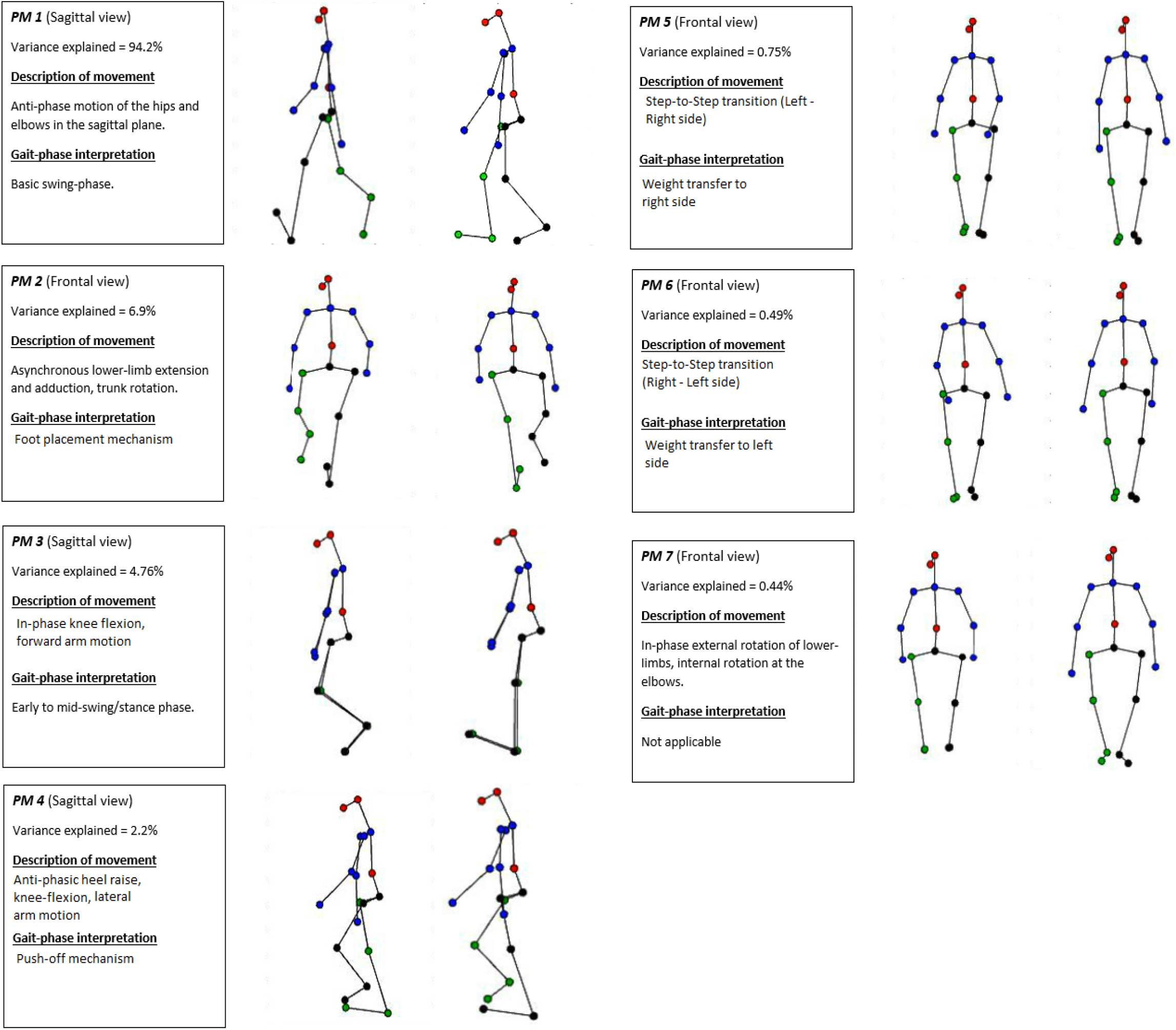
Summaries of each of the validated PM_k_ along with their respective graphical representations to the right. An amplification factor of 2 was used to provide clarity to more subtle movements.

### 3.1.2 Identification of optimal models

Table 2 describes the parsimonious equation 4 models found from a stepwise elimination of effect terms and the findings from a subsequent comparison of these models using Anova. Just two PA_k_RMS components were found to have individual relevance in explaining the variance in DP, that of PA2RMS and PA_4_RMS. A random-participants intercepts term ((1 |Participant)) and a random-participants intercepts and slopes term ((PA_4_RMS |Participant)) were also included in these equation 4 models. The accompanying random-effect term to the PA_4_RMS model indicates that the effect of PA_4_RMS differs significantly in starting value and in the rate of change (i.e. slope) between-participants. The PA_2_RMS model was more parsimonious (degrees of freedom=4) but demonstrated a slightly less goodness-of-fit than the PA_4_RMS model. This difference was not significant however, as indicated by the chi-square statistic (p>0.05). Table 3 describes the findings from a similar procedure with equation 5 models. PM_2_RMS – PM_6_RMS found individual relevance in moderating the relationship between lower-limb DP and the smoothness of arm-swing elevation (DP × *NJI*). All of these models were accompanied by a random-participants intercepts term ((1 |Participant)), indicating within-participant clustering was a predominant level II effect. The PM_2_RMS equation 5 model was used as the baseline model for the Anova. Three equation 5 models were found to be significantly different from this baseline model in terms of goodness-of-fit, with PM_6_RMS conveying the best-fit.

**Table 2:** Output from a comparison of identified equation 4 models using Anova. The PA_4_RMS model was found to be the best fit however insignificantly.

| Goodness-of-Fit indices | DP ~ PA<sub>2</sub>RMS + (1 \|Participant) | DP ~ PA<sub>4</sub>RMS + (PA<sub>4</sub>RMS \|Participant) |
| --- | --- | --- |
| Degrees of freedom | 4 | 6 |
| AIC | 192.5 | 191.0 |
| BIC | 199.2 | 201.0 |
| Log-likelihood | -92.3 | -89.5 |
| Deviance | 184.5 | 179.0 |
| Chi-square (p-value) | - | 5.5 (p>0.05) |

**Table 3:**
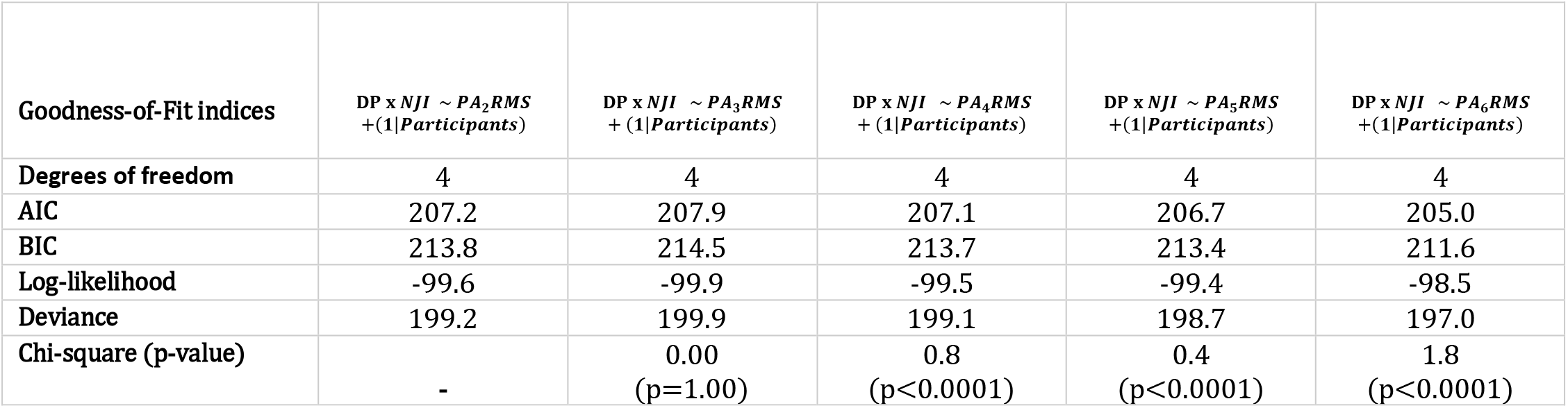
Output from a comparison of identified equation 5 models using Anova. The PA_6_RMSmodel was found to be the best fit.

| Goodness-of-Fit indices | DP x NJI ~ PA<sub>2</sub>RMS<br>+ (1 \|Participants) | DP x NJI ~ PA<sub>3</sub>RMS<br>+ (1 \|Participants) | DP x NJI ~ PA<sub>4</sub>RMS<br>+ (1 \|Participants) | DP x NJI ~ PA<sub>5</sub>RMS<br>+ (1 \|Participants) | DP x NJI ~ PA<sub>6</sub>RMS<br>+ (1 \|Participants) |
| --- | --- | --- | --- | --- | --- |
| Degrees of freedom | 4 | 4 | 4 | 4 | 4 |
| AIC | 207.2 | 207.9 | 207.1 | 206.7 | 205.0 |
| BIC | 213.8 | 214.5 | 213.7 | 213.4 | 211.6 |
| Log-likelihood | -99.6 | -99.9 | -99.5 | -99.4 | -98.5 |
| Deviance | 199.2 | 199.9 | 199.1 | 198.7 | 197.0 |
| Chi-square (p-value) | - | 0.00<br>(p=1.00) | 0.8<br>(p<0.0001) | 0.4<br>(p<0.0001) | 1.8<br>(p<0.0001) |

### 3.1.3 Final mixed-effects models

Table 4 details the regression output from the final, optimal models for both equation 4 & 5 found from the stepwise elimination procedure. Both of the models fixed-effects reached a Bonferroni corrected significance level of p<0.007. PA_4_RMS, the fixed-effect term in the equation 4 model, had a negative effect on DP (β= −2.18±0.51) and explained 37% of the variance in DP (McFadden’s Pseudo R^2^=0.37). With the addition of the random-participants intercepts and slopes term, the equation 4 model explained 70% of the total variance in DP (McFadden’s Pseudo R^2^=0.37). The variance within the data that could be explained by within-participant clustering was σ^2^=3.08 in the equation 4 model and to a lesser extent in the equation 5 model (σ^2^=2.27). Between-cluster variance was also predominant in the data as the random-effect residual variance was σ^2^=3.84 in the equation 4 model and σ^2^=5.32 in the equation 5 model. The inter-class correlation within both models was moderate with an ICC of 0.44 and 0.34 for equation 4 & 5 respectively, indicating the presence of clustering in the data and further demonstrating the necessity for a mixed-effects regression approach. Random-slopes for participants were also included in the equation 4 model and this term captured 1.37 of the variances.

**Table 4:** Optimal models found for equation 4 & 5 (*FE= Fixed effect, RE= Random effect).

| Regression output | **Equation 4**<br>DP ~ PA<sub>k</sub>RMS + (PA<sub>k</sub>RMS \|Participant) | **Equation 5**<br>DP x Jerky ~ PA<sub>k</sub>RMS + (1\|Participant) |
| --- | --- | --- |
| FE $\beta$ Coefficient * | -2.18 | -1.02 |
| FE Standard error* | 0.51 | 0.33 |
| P-value | <0.005 | <0.005 |
| RE-Participant intercepts variance* | 3.08 | 2.27 |
| RE-Participant slopes variance* | 1.37 | - |
| RE-residual variance* | 3.84 | 5.32 |
| Inter-class coefficient (ICC) | 0.44 | 0.34 |
| Pseudo-R<sup>2</sup> (Fixed-effects only) | 0.37 | 0.17 |
| Pseudo-R<sup>2</sup> (Total) | 0.70 | 0.45 |
| Restricted-Maximum Likelihood | 178.3 | 196.3 |

## 4.1 Discussion

This study sought to investigate the role of the neuromuscular system in controlling key movement synergies and how they relate to gait adaptability in healthy adults across various speeds. For this investigation, a coordination metric was devised in the form of deviation phase (DP) that would indicate the level of gait adaptations required in the lower-limbs during a trial of perturbed-walking on a treadmill. The association between this metric and PCA derived measures (PA_k_RMS) representing the magnitude of neuromuscular control on normal-walking movement synergies was then determined using a linear mixed-effects model. The hypothesis that specific PA_k_RMS would have a negative effect on DP was substantiated. As a secondary aim that extended this finding, the current study investigated the moderating effect of PA_k_RMS on the relationship between DP and the smoothness of arm-swing elevation (DP × NJI) during perturbed-walking. The hypothesis that PA_k_RMS would demonstrate a negative effect on this relationship was also substantiated. A stepwise approach was undertaken for both aims as it was foreseeable that a number of movement synergies could be related to DP and (DP × NJI) as differences between participants in one movement synergy feed into differences in others across the gait-cycle (i.e. reciprocal compensation) (Latash, Scholz, & Schöner, 2002). The identification of an optimal model allowed for the main source for these differences to be determined. A number of the identified synergies have been described in relevant research including the push-off and foot placement mechanisms (Fettrow *et al.*, 2019; Reimann *et al.*, 2018), demonstrating the utility of this approach. The current study found the push-off mechanism to be optimal in explaining variance in DP while synergies related to the transfer of weight to the left side were also most relevant for the DP × NJI interaction.

### 4.1.1 Aim 1

The push-off mechanism has been observed in previous research where the modulation of ankle plantar-flexion during a perturbation allowed for the modulation of other gait parameters (i.e. step length and width) (Fettrow *et al.*, 2019; Reimann *et al.*, 2018). Participants in the current study exhibiting a greater control during push-off prior to the slip-event were perhaps more capable of recovering posture by avoiding an excessive foot-floor angle post-slip, a gait parameter associated with slip severity (Moyer *et al.*, 2006). Nazifi and colleagues (2019) found significant associations between slip severity and baseline muscle synergy patterns among healthy adults. Less severe slippers demonstrated higher activation of the medial-hamstring at heel-strike and of the vastus-lateralis following heel-strike allowing for more efficient weight transfer than their less stable counterparts. The findings of the current study mirror this in that the foot placement mechanism was related to DP during a trial of perturbed-walking. Nazifi et al. (2019) focused solely on post-slip synergies and so understandably, the proactive role of the push-off mechanism for gait adaptations was not elicited.

Robert et al. (2009) noted the closed-loop control of gait during the double-support phase and how the CNS appears to use this phase to correct whole-body angular momentum while swing-phase plays a reactive, stabilising role in terms of stride-to-stride reproducibility. It is therefore unsurprising to find a movement synergy identified during the double-support phase of normal-walking was most relevant to DP as this gait-phase plays a proactive role in dynamic balance control. Interestingly, Naizifi et al (2020) was unable to find differences between mild- and severe-slippers in terms of baseline sagittal plane angular momentum, centre-of-mass height, single/double-support duration or upper-body extremity kinematics but did find differences between groups during perturbed-walking. The results of the current study suggest PCA derived movement synergies may be useful in identifying such differences relevant to slip severity during normal-walking trials. Due to the significant random-participant slopes term within the equation 4 model, the current study’s results also suggest that the closed-loop control represented in PA_4_RMS may differ in its effect on gait adaptability significantly across walking-speeds. As this random-effects term was participant-specific rather than trial-specific, this relationship may potentially manifest as a function of the participants preferred walking-speed (H. G. Kang & Dingwell, 2008). Further research is required however to substantiate this claim.

### 4.1.2 Aim 2

Three PA_k_RMS were identified to contribute a moderating effect, that of PA_4_RMS, PA_5_RMS and PA_6_RMS which were interpreted to represent the push-off mechanism and weight-transfer to the right and left sides respectively. Moreover, PA_6_RMS was identified to be the most optimal synergy in explaining the variance in DP × NJI, indicating the central role of this synergy in differentiating healthy participants. Inkol & colleagues (2019) established the role of the upper-limbs in adjusting the centre-of-mass position when a perturbation is experienced. These arm-motion strategies were said to include increased elevation and lateral orientation and depended on the side of the perturbation experienced. They found the frontal plane position of the trunk at heel-strike to be related to perturbed-side arm elevation in that with increased trunk lean towards the trailing limb-side, the CNS intervened by adjusting the elevation of the perturbed-side arm to improve the centre-of-mass position. The current study would support these findings with relevant movement synergies identified during the double-support and weight-transfer phases of gait.

In patient populations where reduced gait stability is present (e.g. stroke), the transfer of weight is typically difficult as strength deficits among extensor muscles make this transfer less efficient (Hsiao *et al.*, 2017). Within healthy populations however, where muscle strength should be sufficient to support normal gait, more subtle differences may still be present that become relevant when compared across healthy participants. Promsri and colleagues investigated the effect of leg dominance on balance control during over-ground single leg-stance (Promsri, Haid, & Federolf, 2018) and also single-legged stance on a multiaxial unstable board (Promsri *et al.*, 2020). Sufficient evidence was found to warrant the consideration of leg dominance in clinical testing and balance training. In the current study, it is interesting that PA_6_RMS, representing right-to-left side weight transfer, was the most relevant movement synergy in this sample despite explaining less variance than the opposite side (PA_5_RMS). Assuming this random sample of healthy adults who were invited to take part in this study were in fact randomly recruited, then it is reasonable to propose that the majority of participants were right-side dominant (Y. Kang & Harris, 2000; Peters & Durding, 1979). Combining this insight with that of the aforementioned observations made by Inkol & colleagues (2019), we propose that asymmetry in limb control between the dominant and non-dominant sides was relevant in differentiating healthy adults in terms of gait adaptability in this study. The moderating effect of movement synergies on coupled asymmetries between the upper- and lower-limbs has been found in recent research and may have significant influence on gait performance and risk for injury (Ó’ Reilly, 2020). The influence of these inherent asymmetries on gait stability has not been investigated in the research thus far. We therefore concur with the aforementioned studies related to leg dominance that this phenomenon should be considered in balance testing and clinical evaluations.

## 5.1 Limitations

Healthy adults in this study were subjected to perturbations during treadmill walking and therefore the findings cannot be generalized directly to over-ground walking. Results should be interpreted with the limitations of a relatively small sample size in mind.

## 6.1 Conclusion

This study identified movement synergies sourced from normal-walking trials whose magnitude of neuromuscular control had significant negative effects on lower-limb coordination variability during perturbed-walking at various walking-speeds. Relevant movement synergies were also identified that moderated the relationship between lower-limb coordination variability and the smoothness of arm-swing motions, thus indicating the utility of this approach in differentiating healthy adults in terms of gait adaptability. Future research should investigate further the change in control of these movement synergies across walking-speeds, the influence of leg dominance on gait adaptability and the utility of this analytical approach in patient populations.

## Abbreviations

CNS: Central nervous system
CRP: Continuous relative phase
DP: Deviation phase
ICC: Inter-class correlation
NJI: Normalised Jerk Index
PA_k_: Principal accelerations
PCA: Principal component analysis
PM_k_: Principal Movements
PV_k_: Principal velocities
RMS: Root-mean-square

## 8.1 Author contributions

David Ó’Reilly: conceptualization, methodology, software, formal analysis, writing – original draft.

Peter Federolf: resources, writing – review & editing.

